# Directed Conservative Causal Core Gene Networks

**DOI:** 10.1101/271031

**Authors:** Gokmen Altay

**Affiliations:** La Jolla Institute for Allergy and Immunology, 9420 Athena Circle, La Jolla, CA 92037, USA

## Abstract

**Motivation:** Inferring large scale directional networks with higher accuracy has important applications such as gene regulatory network or finance.

**Results:** We modified a well-established conservative causal core network inference algorithm, C3NET, to be able to infer very large scale networks with direction information. This advanced version is called Ac3net. We demonstrate that Ac3net outperforms C3NET and many other popular algorithms when considering the directional interaction information of gene/protein networks. We provide and R package and present performance results that are reproducible via the Supplementary file.

**Availability:** Ac3net is available on CRAN and at github.com/altayg/Ac3net

**Supplementary information:** Supplementary file is available online.

## 1 Introduction

There are many algorithms developed to infer causal large-scale (e.g. >10000 variables) gene regulatory networks with higher accuracy. Conservative Causal Core Network inference algorithm, C3NET (Altay and Emmert-Streib, 2010), is a very simple but an elegant algorithm that outperformed many other popular algorithms such as ARACNE (Margolin, et al., 2006), MRNET (Meyer, et al., 2008), CLR (Faith, et al., 2007) and RN (Butte, et al., 2000). The comparative analysis in (Altay and Emmert-Streib, 2010) was performed on 6 different topologies and more than 1000 different datasets that reflects more generic comparative results. C3NET gave birth to many other algorithms (Altay, et al., 2011; Legeay, et al., 2016; Simoes and Emmert-Streib, 2012) and is utilized not just in the field of genomics but also in finance (Baltakys, et al., 2017). C3NET is two steps, first it eliminates all the statistically non-significant dependency score values (mutual information) of the adjacency matrix, then in the second step it selects the maximum valued partner for each gene in the adjacency matrix. However, since it does not assume direction, the interaction obtained from the maximum valued partner can be in either direction. This is because, C3NET was designed to infer non-directional networks. Therefore, in the **equation 5** in (Altay and Emmert-Streib, 2010), where C3NET was introduced, the algorithm converts the non-symmetric matrix into symmetric and ensures that the network is non-directional. This symmetry forcing step is also available in the mathematical algorithmic description of C3NET in **Algorithm 1** at **number 17** in (Altay and Emmert-Streib, 2010). In this study, we modify the algorithm C3NET to infer directional networks instead of non-directional networks.

## 2 Methods

In order to infer directional networks, we put forward the following null hypothesis:

**H**_0_: A variable in an adjacency matrix is regulated by its maximum scored (e.g. correlated) partner.

The results of the analysis of this document stand as a solid evidence that support the hypothesis. Therefore, based on the assumed null hypothesis in mind, we modified C3NET as follows. The essential and most discriminative difference is that we remove the **equation 5** or the **number 17** of the **Algoritgm 1** in (Altay and Emmert-Streib, 2010) from C3NET and assume the remaining network is directional based on the above defined null hypothesis. Therefore, in this network, an interaction is directed from the maximum valued partner (or partners) to the gene.

Ac3net has also another difference, which should be mentioned though tiny. If a gene has more than one maximum valued partner, then all of the maximum valued partners are included in the network as the regulator of that gene. This inclusion of all the maximum partners is not available in C3NET. Although this point was not discussed in its paper, it takes the first maximum in its R package implementation on CRAN repository (*c3* function). This tiny difference of including all the maximum valued partners instead of just one was realized by the study in (Legeay, et al., 2016), but in that study the direction of the interactions is not specified.

Since the proposed algorithm is essentially different and an **a**dvanced version of the well-established algorithm C3NET, we name it as **A**c3net. With Ac3net, we can infer directional networks at very large scale datasets (e.g. > 20000 variables and 100 samples), which is not practical with Bayesian network inference approaches (Kjaerulff and Madsen, 2008). There are also other differences of Ac3net with compare to C3NET, which are not critical in general as they can be exchanged in the future if better alternatives appear for them, as follows. In C3NET, the default dependency score estimator was the information theoretical estimators, parametric Gaussian estimator or Pearson based Gaussian (PBG), which provides mutual information (MI) values that are always positive and there is no maximum value. Instead, in Ac3net, we use Pearson Correlation Coefficient (PCC) with normalized data (at least Log-2) to estimate the dependency scores between the variables. We observed no performance differences in general between PCC and PBG. In cases of unnormalized data, we use Spearman Correlation Coefficient (SCC). Since we use PCC, the dependency scores have signs that show the type of interaction which is not available in any MI based algorithms like C3NET, ARACNE, MRNET, CLR and RN. Moreover, since PCC has the highest absolute magnitude as 1, we can set an arbitrary PCC threshold above |0.5|. As a moderate PCC significance threshold, we can use |0.75| or with a stricter threshold |0.85| with no additional computational cost to get a significance threshold. As opposed to C3NET algorithm, we also do not use the copula transformation before computing dependency scores. With this study, we also present the R package of the algorithm Ac3net, which is available on CRAN repository. We define the proposed algorithm Ac3net as follows.

### Ac3net algorithm

Step 1. A is adjacency matrix (with optionally PCC values) and the nonsignificant values below a significance cutoff value are set to zero.

Step 2. G_i,j_ is **inferred** as an interaction **if** the absolute maximum non-zero valued partner of the variable (e.g. gene) *i* is the variable *j*. The **direction** of this interaction is from variable *j* to variable *i* (based on H_0_ as defined above). Also, *i* can have more than one maximum valued partner.

The attachment of this paper presents a reproducible comparative analysis study of Ac3net with the mentioned popular algorithms. Here we shortly present its results.

## 3 Results

First of all, we need to define the example dataset we used and also the performance metrics of the analysis along with some nomenclature.

TP: True Positive, the inferred interactions that are also in the reference network. FP: False Positive, the inferred interactions that are not in the reference network. FN: False Negative, the interactions that are in the reference network but were not inferred. TN: the interactions that are not inferred and they are not in the reference network. Accuracy is the main metric we chose to compare algorithms as it contains all the performance scores.

Accuracy = (TP + TN) / (TP + TN + FP + FN)

Precision = TP / (TP + FP)

Recall = TP / (TP + FN)

F-score = 2 x Precision x Recall / (Precision + Recall)

In the Ac3net R package, there is *Ac3net.performance* function that provides all the above performance scores.

The example gene expression dataset we used is the very popular *E.coli* gene expression dataset which was compiled in (Faith, et al., 2007) and it has 1146 genes and 524 samples. This dataset can be easily accessed from the presented Ac3net R package by the function *data(expdata)*. The reference or true network of this dataset was downloaded (Faith, et al., 2007) from RegulonDB database (Gama-Castro, et al., 2016). This dataset is being used intensely as a real gene expression dataset with available reference network by scientific community to be able to evaluate the performance of new algorithms. It was also used in one of DREAM (Dialogue for Reverse Engineering Assessments and Methods) project (dreamchallenges.org) and its conference series (DREAM2, Challenge 5, Genome-Scale Network Inference). In order to get more updated reference network, we recently downloaded gene-gene and transcription factor (TF)-gene interactions databases from RegulonDB database and combined it with the previously available reference network from (Faith, et al., 2007). We filtered the interactions from genes that are not available in the example gene expression dataset. This reference network can be easily accessed with the Ac3net R package using the function *data(truenet)*. There are 3792 true interactions in this reference network.

All the performance analysis presented here, with further details, can be seen and reproduced with the Supplementary File of this paper. In the first analysis, we will compare the inferred directional network of Ac3net with the network obtained by the opposite direction of Ac3net network to see if there is a difference in performance with regard to direction information.

As we see from Table 1, the direction inferred by Ac3net provides dramatically better performances than the opposite. Ac3net inferred directional network provides % 235.7 more precision than the opposite direction. There are 38 TPs that come from the dual interactions that are common TPs in both networks. Removing the common performance effect, the Ac3net directional network, without dual links, gives %2475 TP increase with compare to the opposite direction. Also, the Ac3net directional network, without dual links, provides %34 FP decrease with respect to the opposite direction. These results show that the direction inferred by Ac3net provides significantly better performances. The results stand as a solid evidence that supports the null hypothesis that is the base of the proposed algorithm Ac3net, which allows Ac3net to determine the direction of the inferred interaction.

**Table 1.**
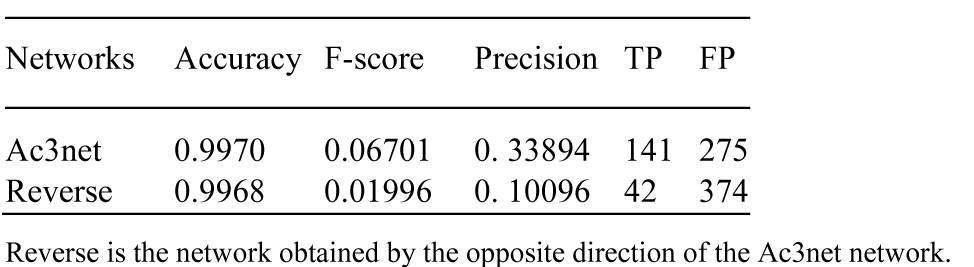
Performances regarding directions of interactions

In the second analysis, we will compare the performance of Ac3net with the other algorithms.

Table 2 shows that when the direction information of the reference network is considered, which is the nature of gene interaction data and correct way, then Ac3net provides the best Accuracy. The difference in the Accuracy results can be seen in more resolution in the Precision scores, where there is a dramatic difference between the results of Ac3net and the other algorithms.

**Table 2.**
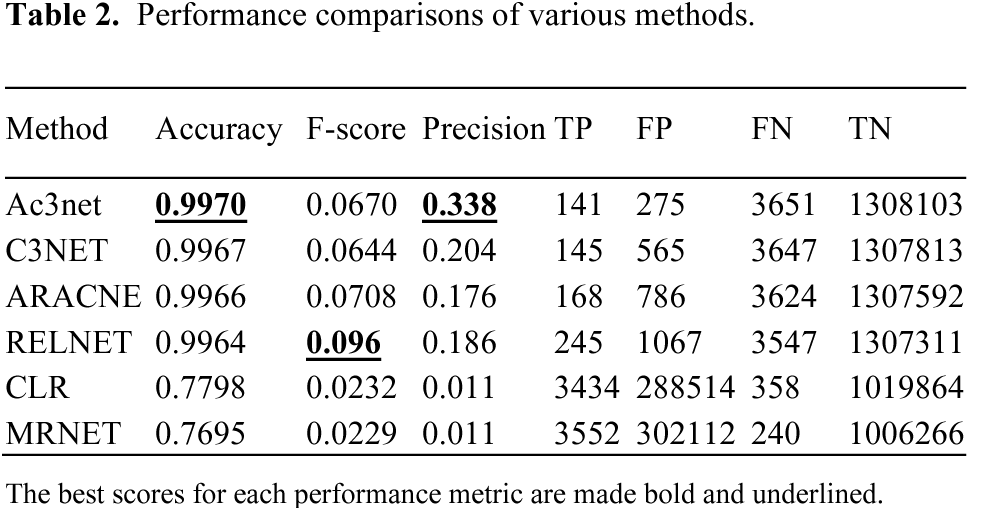
Performance comparisons of various methods.

As a conclusion, we presented an algorithm and its R software package that allow inferring directional networks at very large scale and showed that it provides higher performance than other popular algorithms when directionality is taken into account. As the directionality information mostly exist in real life problems, such as gene regulatory networks, the presented algorithm, Ac3net, might be more useful in applying to the very large scale real data applications.

## 4 Ac3net R package

Ac3net can be accessed from CRAN or github.com/altayg/Ac3net repositories. Assuming the data object has name *data*, the simplest usage of it in R can be as follows: net = Ac3net(data). In this network, regulators are on the first column, targets are on the second column and their dependency scores are on the third column. There are some parameters that might need to be set differently depends on the application. For the details and example usage we suggest reading the Reference Manual of the package and also the Supplementary file of the paper which reproduces all the results of the analysis in this paper.

